# RNA-binding proteins regulate aldosterone homeostasis in human steroidogenic cells

**DOI:** 10.1101/2021.02.19.431050

**Authors:** Rui Fu, Kimberly Wellman, Amber Baldwin, Juilee Rege, Kathryn Walters, Antje Hirsekorn, Kent Riemondy, William Rainey, Neelanjan Mukherjee

## Abstract

Angiotensin II (AngII) binds to the type I angiotensin receptor in the adrenal cortex to initiate a cascade of events leading to the production of aldosterone, a master regulator of blood pressure. Despite extensive characterization of the transcriptional and enzymatic control of adrenocortical steroidogenesis, there are still major gaps in our knowledge related to precise regulation of AII-induced gene expression kinetics. Specifically, we do not know the regulatory contribution of RNA-binding proteins (RBPs) and RNA decay, which can control the timing of stimulus-induced gene expression. To investigate this question, we performed a high-resolution RNA-seq time course of the AngII stimulation response and 4-thiouridine pulse labeling in a steroidogenic human cell line (H295R). We identified twelve temporally distinct gene expression responses that contained mRNA encoding proteins known to be important for various steps of aldosterone production, such as cAMP signaling components and steroidogenic enzymes. AngII response kinetics for many of these mRNAs revealed a coordinated increase in both synthesis and decay. These findings were validated in primary human adrenocortical cells stimulated ex vivo with AngII. Using a candidate siRNA screen, we identified a subset of RNA-binding protein and RNA decay factors that activate or repress AngII-stimulated aldosterone production. Among the repressors of aldosterone were BTG2, which promotes deadenylation and global RNA decay. BTG2 was induced in response to AngII stimulation and promoted the repression of mRNAs encoding pro-steroidogenic factors indicating the existence of an incoherent feedforward loop controlling aldosterone homeostasis. Together, these data support a model in which coordinated increases in transcription and regulated RNA decay facilitates the major transcriptomic changes required to implement a pro-steroidogenic gene expression program that is temporally restricted to prevent aldosterone overproduction.

## INTRODUCTION

Angiotensin II (AngII) is a potent octapeptide hormone that binds to angiotensin II receptor type 1 (AGTR1) in adrenal zona glomerulosa cells to stimulate the production of the mineralocorticoid aldosterone from cholesterol. The primary function of aldosterone is to regulate blood pressure and volume through the control of renal salt excretion [cite]. Additional functions of aldosterone discovered more recently include directly mediating cardiac and vascular remodeling [cite]. Aldosterone is a small, non-polar, and hydrophobic steroid hormone that passes through the cell membrane upon production. Given the lack of intracellular aldosterone, it must be produced de novo in response to steroidogenic stimuli. Thus, it is critical to tightly control the timing and amount of de novo aldosterone production. Pathological aldosterone excess, such as in primary aldosteronism, leads to hypertension and increased cardiovascular risk in humans (Monticone *et al*, 2018; Nanba *et al*, 2019; Brown *et al*, 2020). Although the stimulatory effect of AngII upon aldosterone secretion has been extensively studied, the temporal regulation and resolution of the AngII response remains poorly understood and critical for maintaining aldosterone homeostasis.

The temporal coordination of AngII-stimulated aldosterone biosynthesis is described in two stages: an “early regulatory step” (minutes) and a “late regulatory step” (hours) (Nogueira *et al*, 2009; Hattangady *et al*, 2012). The early regulatory step is characterized by rapid signaling pathways promoting cholesterol mobilization. A key part of the early step is inducing the expression of Steroidogenic Acute Regulatory Protein (STAR) that transports cholesterol from the outer to inner mitochondrial membrane allowing for conversion by CYP11A1 into the pregnenolone, which is a precursor for the production of all steroid hormones including aldosterone. The late regulatory step is characterized by inducing the expression of steroid biosynthesis enzymes such as aldosterone synthase (CYP11B2). Although the temporal order of the AngII gene expression response underlies both regulatory steps there has not been an examination of steroidogenic transcriptome dynamics with sufficient temporal resolution to systematically identify distinct response kinetics.

There are two prototypical examples of coordinated temporal response utilize very different regulatory strategies. One example utilizes a cascade of transcription factor activity to generate waves of temporal expression patterns. Transcriptional cascades have been observed in the yeast cell cycle and innate immune responses (Luscombe *et al*, 2004; Smale, 2012). The other example relies on intrinsic differences in the stability between induced transcripts to generate sequential response response patterns.Specifically, the time to peak response is inversely correlated with the stability of the transcript, and thus unstable RNAs will exhibit more rapid induction and return to baseline than stable RNAs (Palumbo *et al*, 2015). This post-transcriptionally regulated model of temporal coordination of response kinetics has been observed largely in immune and inflammation responses (Hao & Baltimore, 2009; Elkon *et al*, 2010; Rabani *et al*, 2011). Interestingly, increasing the decay rate of transcriptionally induced RNAs results in a kinetic response with a quicker time to peak expression and return to baseline. This strategy of increasing synthesis and decay has been observed in the yeast oxidative stress response and immune activation of mouse dendritic cells (Shalem *et al*, 2008; Rabani *et al*, 2014). A lack of response resolution in both of those biological systems could clearly lead to deleterious effects for the cell and organism, much like the uncontrolled resolution of aldosterone production.

RNA-binding proteins (RBPs) are a class of trans-acting regulatory factors that control all aspects of RNA metabolism (Gerstberger *et al*, 2014). Many RBPs have been shown to control the stability of mRNAs through interactions with cis-regulatory elements in the 3’ UTR of their target RNAs (Keene, 2007). RBPs binding to AU-rich elements are one of the most well-characterized examples for the regulation of RNA stability and have been associated with controlling the temporal order and shape of immune response kinetics (Elkon *et al*, 2010; Hao & Baltimore, 2009; Rabani *et al*, 2014). General regulators of RNA stability, such as factors that modify deadenylation, can also control the temporal response kinetics of induced RNAs and may do so in cardiac hypertrophy (Mauxion *et al*, 2008; Stupfler *et al*, 2016; Masumura *et al*, 2016). In spite of the role of RBPs in the regulation of decay, expression kinetics, and many other post-transcriptional processes, their role in AngII-mediated aldosterone production is poorly characterized.

To understand the molecular basis and role of RNA regulation in the kinetics of the AngII-aldosterone gene regulatory program, we measured transcriptome dynamics at high-resolution temporal by RNA-seq in a steroidogenic human cell line (H295R) stimulated with AngII. While transcriptional control was clearly prominent, RNA decay rates determined the time to maximal change in expression. Furthermore, we observed AngII-dependent increases in RNA decay for many transcriptionally induced RNAs, including those encoding key pro-steroidogenic proteins. These findings were validated in primary human adrenocortical cells stimulated ex vivo. We identify RBPs and RNA decay factors that either activate or repress aldosterone production. Finally, we find that AngII-induced RNA decay factor BTG2 represses aldosterone production and promotes the clearance of steroidogenic RNAs. Together, these data support a model in which coordinated increases in transcription and RNA decay facilitates the major transcriptomic remodeling required to implement a pro-steroidogenic gene expression program that is temporally restricted to prevent aldosterone overproduction.

## RESULTS

### Identification of genes with temporally distinct AngII-response kinetics in H295R cells

To delineate the temporal coordination of the AngII-induced gene expression response, we performed a high-resolution (twelve time points in duplicate) AngII stimulation RNA-seq time course in H295R cells (Figure 1A). Rather rathan polyA-enrichment, we performed rRNA depletion to enable measurement of both precursor and mature RNAs. The data were high quality as samples clustered first by replicates (r ∼ 0.99) and then time post-stimulation (Supplemental Figure 1A, 1B). First, we focused on changes in mature RNA (sum of all mature transcripts per gene, Supplemental Table 1). PCA analysis clearly indicated differences in gene expression as early as 15 minutes with the largest remodeling of the transcriptome occurring ∼6-8 hrs post-stimulation (Figure 1B). Furthermore, the return towards the unstimulated state by 24 hrs suggests that this time course captures most of the AngII-induced changes in RNA levels in this system. Next, we determined the genes exhibiting statistically significant changes (FDR <= .001 and > 2-fold change) in mature RNA expression at any time point of AngII treatment versus unstimulated (n=2417). The changes in expression measured by RNA-seq were recapitulated (R > .95) by qRT-PCR for a subset of differentially expressed genes (Supplemental Figure 1C). Differentially expressed coding and long-noncoding RNAs both exhibited the largest change in expression between 3 and 8 hours (Figure 1C). Intriguingly, snRNAs appeared to be upregulated rapidly, peaking around 2 hrs before returning to baseline.

**Figure 1.**
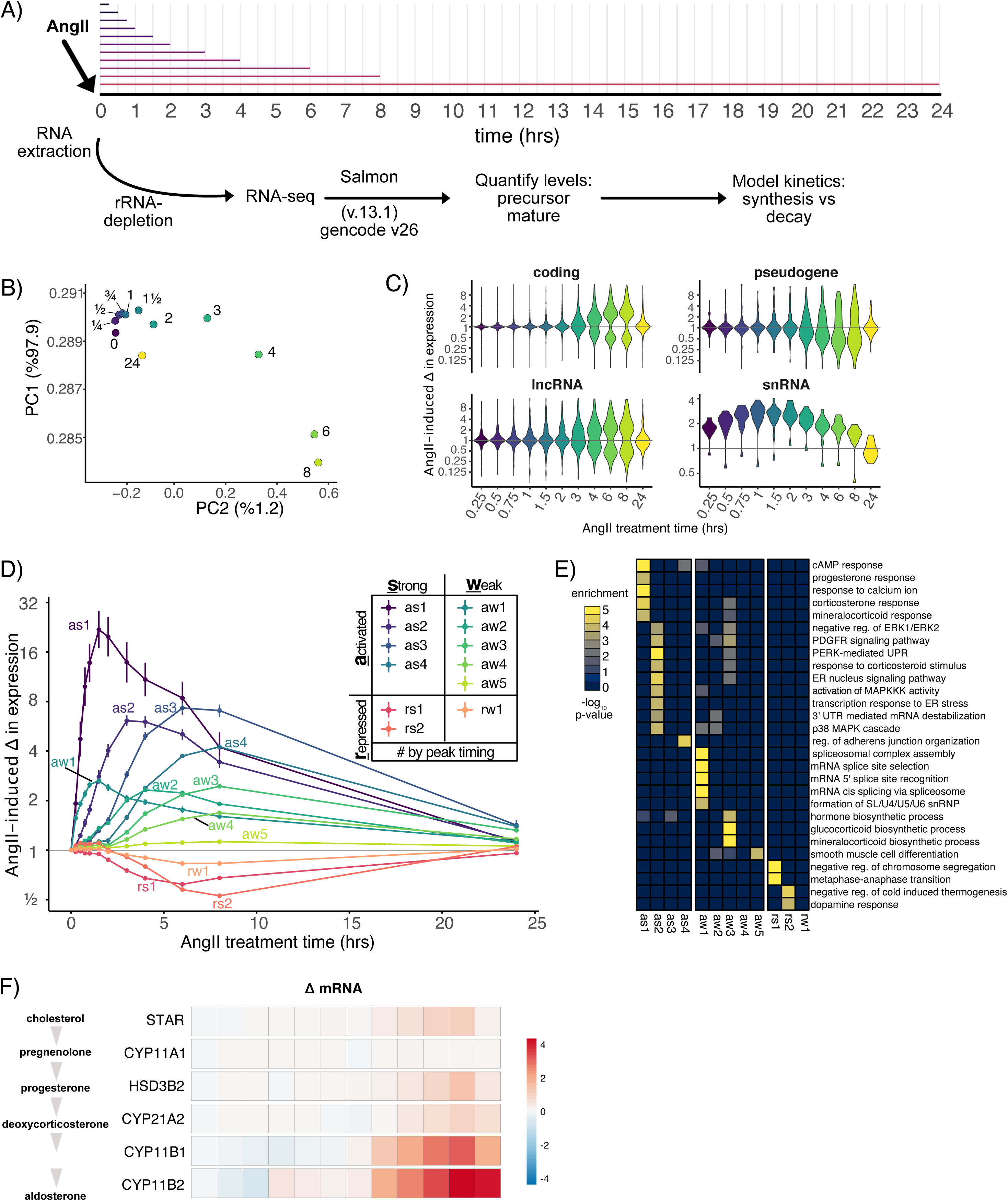
High-resolution AngII-response kinetics in H295R cells. A) Cartoon of AngII treatment time course RNA-sequencing experiment. Total RNA is depleted for rRNA and precursor and mature RNA expression levels are calculated allowing modeling of synthesis and decay. B) Scatter plot of the first two principal components calculated on a matrix of the mean expression level for each expressed gene at each time point (colors) after AngII stimulation. C) Violin plot of the distribution of genes exhibiting statistically significant AngII-induced changes in expression per time point and category. D) Line plot depicting the average change in expression (error bar represents standard error) relative to unstimulated cells for each group of genes with similar response kinetics determined by k-means clustering. Each color represents a specific group of genes with the name defined by the direction, shape, and timing of the AngII-induced response. E) Heatmap of selected gene sets (Molecular Function ontology) that are significantly enriched in the AngII response groups. F) Heatmap of log2-based fold changes in pre-mRNA and mature mRNA levels upon AngII stimulation compared to unstimulated cells for mRNAs encoding key steroidogenic enzymes.

To identify genes with similar temporal changes in expression, we performed K-means clustering on the mature expression level changes for the differentially expressed genes. This revealed 12 groups of genes with temporally distinct AngII-response kinetics (Figure 1D and Supplemental Figure 1D, Supplemental Table 2). The groups were labeled based on the direction (activated/repressed), strength (strong/weak), and timing of expression changes. For example, RNAs belonging to as1 (activated strong 1), exhibited an average maximal activation of ∼20-fold peaking around 1 to 2 hours, while RNAs in as4 (activated strong 4) exhibited an average maximal activation of ∼4-fold peaking around 7 hrs. We asked whether specific gene biotypes were enriched in specific clusters. The as1 RNAs encoded proteins involved in the calcium and cyclic AMP signaling pathway that are activated during steroidogenesis (Figure 1E and supplemental Figure 1E), including NR4A1 and NR4A2, which are known transcriptional regulators of enzymes controlling aldosterone production (Romero *et al*, 2004; Bassett *et al*, 2004). Unexpectedly, we found a very strong enrichment for snRNAs, and particularly U1 snRNA, in aw1 (Supplemental Figure 1F). We observed a strong enrichment in as2 for mRNAs encoding proteins regulating the unfolded protein response. RNA-binding proteins that control transcript-specific RNA decay were enriched in as2 and aw2. The aw3 and as3 groups were enriched for mRNAs encoding steroid biosynthesis proteins. The induction of mRNAs encoding steroidogenic enzymes required for the conversion of cholesterol to aldosterone was consistent with previous studies validating the quality of the data (Figure 1F) (Romero *et al*, 2007; Wang *et al*, 2012). Altogether, our analysis demonstrated that AngII response kinetics exhibited temporal coordination of functionally related genes.

### RNA decay controls AngII response kinetics

Simulations and previous studies in other stimulation-response paradigms have shown that the initial RNA decay rates in the unstimulated cells govern the time to peak expression response and thus the shape of the expression response (Rabani *et al*, 2011; Palumbo *et al*, 2015). Importantly, this phenomenon assumes constant decay rates throughout the response. As a starting point, we quantified the time to maximal expression change for each gene (see methods), which increased concordantly with peak time assigned to each group (Figure 2A, left). For example, genes in as1 (median 1.8 hrs) reached their peak expression 5.7 times faster than those in as4 (median 10.9 hrs). Next to test if RNA decay rates governed maximal response kinetics in our system, we measured decay rates in unstimulated H295R cells using RNA metabolic labeling as we and others have done previously (Mukherjee *et al*, 2017; Milek *et al*, 2017). Cells were pulsed with 4-thiouridine (4sU) for 20 minutes and both total RNA and 4sU-labelled RNA were subject to rRNA depletion and RNA-sequencing, and synthesis, processing, and decay rates were calculated using INSPEcT for 11,668 genes in unstimulated H295R cells (Supplemental Figure 2A-C, Supplemental Table 3) (de Pretis *et al*, 2015). Our decay rates were consistent with established expectations for stable and unstable mRNAs confirming the validity of our measurements. For example, many transcripts encoding ribosomal proteins had low decay rates, while transcripts encoding immediate early genes had high decay rate (Supplemental Figure 2D). We found higher decay rates for temporal groups that peaked earlier post AngII stimulation (Figure 2B, right). For example, RNAs in as1 had a ∼4.5x higher decay rate than RNAs in as4. Indeed, we observed an inverse relationship between the median decay rate and the median time to maximal induction for both strongly and weakly activated genes (Figure 2B). These results demonstrated that initial RNA decay rates in unstimulated cells play an important role in shaping AngII-response kinetics.

**Figure 2.**
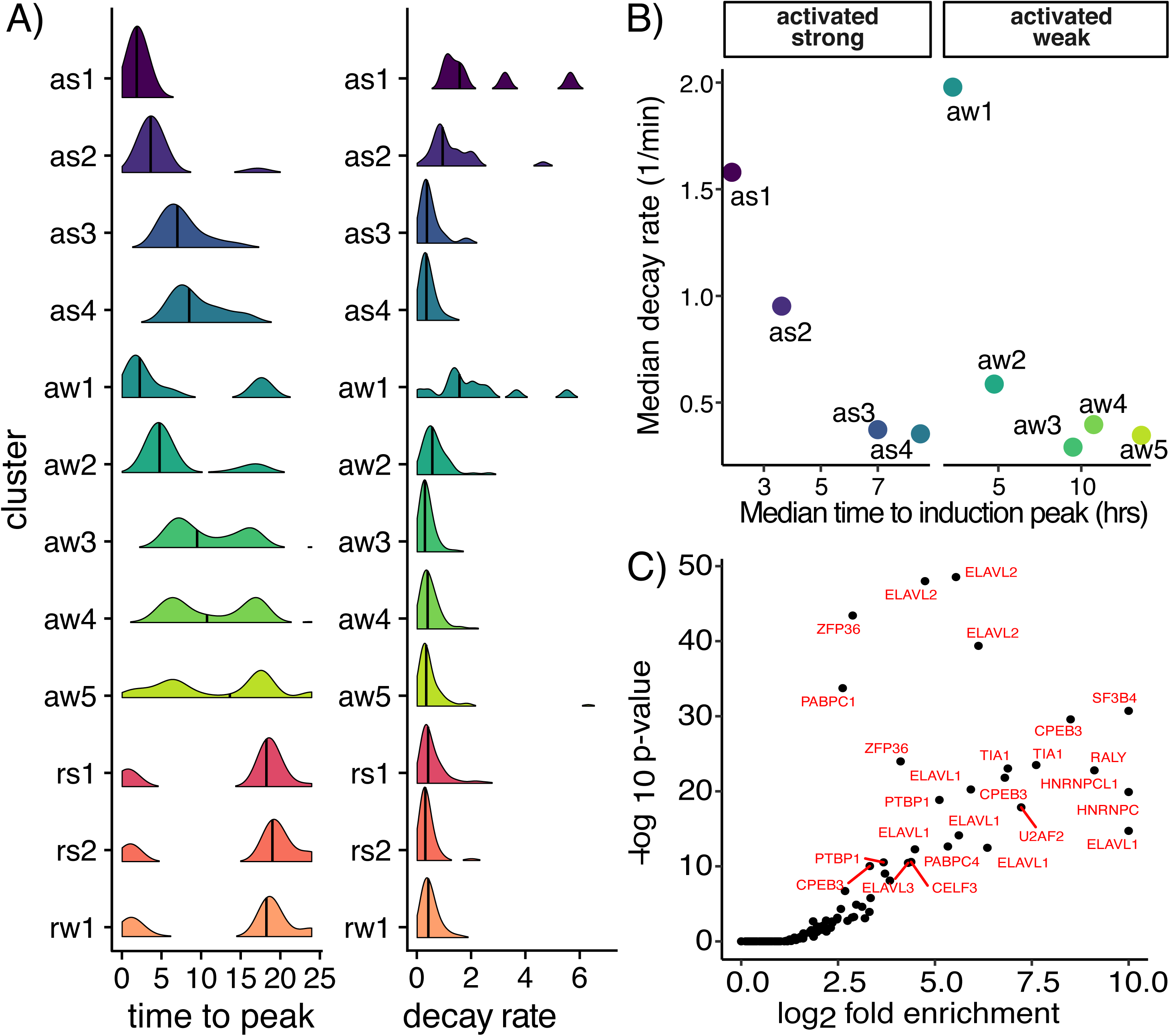
The influence of RNA decay on AngII response kinetics. A) Distribution of time to peak response for all RNAs per cluster (left). Distribution of decay rates in unstimulated cells for all RNAs per cluster determined from 4sU-labeling experiments (right). B) Scatter plot of the average decay rate and peakiness for activated RNAs. C) Scatter plot of the p-value and enrichment of RBP motifs from in the top 500 7mers found in the 3’ UTR of RNAs with high decay rates in unstimulated cells.

Next, we ascertained putative regulatory mechanisms controlling RNA decay rates in steroidogenic cells. To identify possible RBPs or miRNAs regulating stability, we ranked mRNAs by decay rates and used cWords to identify 7-mers over-represented in the 3’ UTR sequence of unstable and stable RNAs (Rasmussen *et al*, 2013). Most of the significantly enriched motifs were associated with unstable RNAs versus stable RNAs. Therefore, we scored the top 500 7mers associated with instability for matches to RBP recognition motifs from external databases. These analyses revealed a striking enrichment for RBPs binding AU-rich elements (AREs), which are key determinants of RNA stability (Figure 2C). We also observed enrichment for RBPs regulating splicing and export, such as U2AF2 and RALY. These data identify putative RBPs and RNA regulatory elements that regulate RNA stability in unstimulated H295R cells. Assuming constant RNA decay rates throughout the AngII-response, higher decay rates in the unstimulated cells inversely correlate with time to peak expression in response to AngII stimulation (Figure 2B. Therefore, RBPs that control the RNA decay rates in unstimulated cells are important regulatory factors controlling AngII response kinetics and ultimately steroidogenesis.

### Increased RNA decay during steroidogenesis

Having established the importance of initial RNA decay rates on AngII response kinetics, we next asked if there was evidence for changes in decay rate during the steroidogenic response. We compared changes in pre-mRNA versus mRNA levels to determine whether gene expression changes were better explained by transcriptional or post-transcriptional regulation. Specifically, we utilized reads aligning to introns versus exons to determine AngII-induced changes in levels of pre-mRNA and mRNA, respectively (Gaidatzis *et al*, 2015; Alkallas *et al*, 2017; Mukherjee *et al*, 2017, 2019). We added the entire precursor RNA sequence and treated it as another transcript isoform to individually quantify precursor (Supplemental Table 4) and mature transcript levels using Salmon (Patro *et al*, 2017). As a first pass, we calculated the correlation between the changes in pre-mRNA and mRNA levels of genes encoding steroidogenic enzymes required for the conversion of cholesterol to aldosterone and found examples that are largely explained by changes in transcriptional regulation (high correlation), as well as examples indicative of changes in post-transcriptional regulation (low correlation) (Figure 3A). The same analysis was performed on all ∼1500 differentially expressed genes with introns and sufficient intronic read support, and revealed a broad distribution of correlation coefficients (Figure 3B). Each temporal group had many genes with low or negative correlation, indicating that changes in transcriptional regulation alone cannot explain the mature RNA changes. We also performed cross-correlation to account for processing delays and observed similar results (Supplemental Figure 3A). These results suggested that many genes were experiencing stabilization or destabilization in response to AngII stimulation.

**Figure 3.**
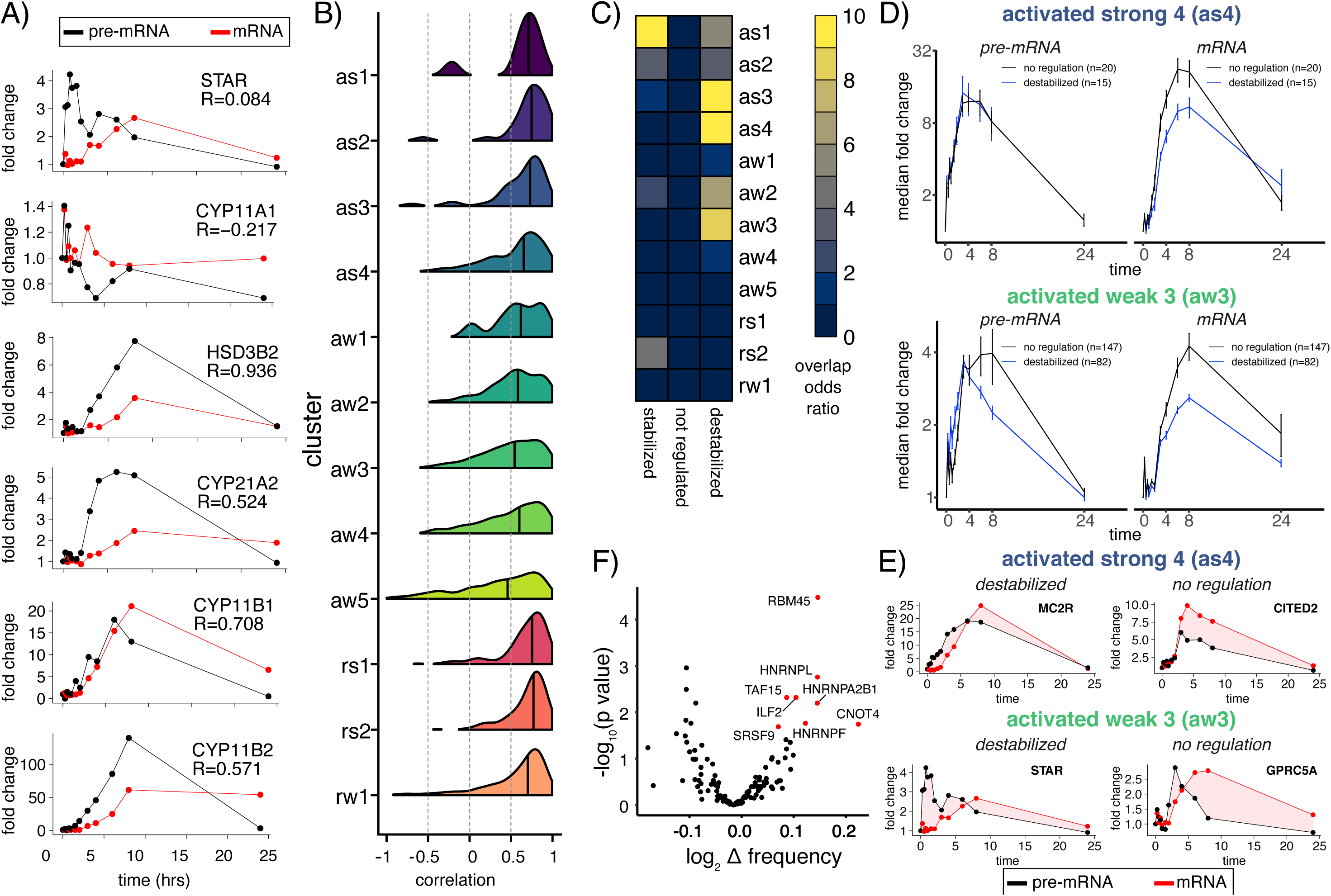
Dynamic changes in RNA decay during steroidogenesis. A) Line plot of fold change in pre-mRNA (black) and mRNA (red) expression for mRNAs encoding enzymes responsible for conversion of cholesterol to aldosterone (top-to-bottom). Pearson correlation coefficient between changes pre-mRNA and mRNA for each gene (top right). B) Distribution of Pearson correlation coefficient between pre-mRNA and mRNA changes for each gene across all time points for each response cluster. C) Heatmap of the odds ratio for the overlap between response cluster membership and RNAs exhibiting evidence for stabilization, destabilization, or neither as determined by EISA (yellow indicates more overlap). D) Line plot of median fold change in pre-mRNA (left) and mRNA (right) expression for genes in as4 (top) or aw3 (below) with either evidence for destabilization (blue) or no change in post-transcriptional regulation (black). Error bar represents standard error. E) Line plot of mean fold change in pre-mRNA (black) and mRNA (red) expression for example genes from as3 (top) or aw3 (below) and evidence for destabilization (left) or no difference in decay rates (right). F) Scatter plot of the p-value and enrichment of RBP motifs in 5mers within the 3’ UTR of destabilized mRNAs in aw3 versus 3’ UTRs of non-differentially expressed mRNAs.

To test the extent and direction of changes in post-transcriptional regulation, we performed exon-intron-split analysis (EISA) using precursor and mature RNA estimates comparing all AngII-stimulated samples to the unstimulated baseline (see methods, Supplemental Table 5). Filtering of the initial 2417 differentially expressed genes for the presence of introns and sufficient intronic read coverage resulted in 1194 genes that were analyzed. Among these genes a large majority (84% n=223) showed evidence of mRNA destabilization. Destabilized transcripts were significantly overrepresented in activated genes that had peak expression between 4 and 8 hours post AngII stimulation (as3, as4, aw2, aw3) (Figure 3C). Next, we examined AngII-induced genes with evidence for destabilization focusing on as4 and aw3, which contained many steroidogenic genes (Figure 1E). The amplitude of AngII-induced changes in precursor RNA were similar between genes with (blue lines) and without evidence for destabilization (black lines), albeit the precursor RNAs for destabilized genes tend to peak earlier (Figure 3D, left). However, the changes in the mature RNA of destabilized genes (blue lines) were clearly suppressed relative to genes without evidence for post-transcriptional regulation (black lines) (Figure 3D, right). Included among the destabilized mRNAs were proteins encoding STAR, which is required for the proper and timely enzymatic conversion of cholesterol to steroid hormones, and MC2R (Lin *et al*, 1995), which is the ACTH receptor known to be induced by AngII stimulation (Lebrethon *et al*, 1994; Parmar *et al*, 2008) (Figure 3E). We identified potential RBPs responsible for the increased decay by searching for RBP binding motifs over-represented in the 3’ UTR of destabilized mRNAs belonging to aw3 and other clusters (Figure 3F) (Supplemental Figure 3B) enriched for destabilized mRNAs. Amongst these RBPs are known regulators of RNA decay ILF2, CNOT4, HNRNPA2B1, and HNRNPL (Hui *et al*, 2003; Han *et al*, 2010; Albert *et al*, 2000). Altogether, we have identified hundreds of genes (n = 223) exhibiting coordinate increases in both transcription and RNA decay, including key steroidogenic genes, and putative RBPs regulating AngII-induced post-transcriptional regulatory dynamics.

### Ex vivo steroidogenic response in primary human adrenocortical cells

Although the expression dynamics in H295R cells stimulated with AngII suggested a key role for RBP-mediated RNA decay, the extent to whether these expression changes occur in normal human adrenocortical steroidogenesis is unclear. Therefore, we performed RNA-seq in adrenocortical cells isolated from the adrenal gland of a tissue donor and treated ex vivo with either AngII or ACTH or basal media for 3 or 24 hrs in triplicate (Figure 4A, Supplemental Tables 6 and 7) (Xing *et al*, 2010, 2011). We included ACTH, which stimulates the production of cortisol by the zona fasciculata, because H295R cells do not correspond to a specific adrenocortical zone but produce cortisol and androgens in addition to aldosterone when stimulated with AngII (Parmar *et al*, 2008). Replicates were highly correlated and clustered by treatment condition and time (Supplemental Figure 4A). PCA analysis revealed that compared to basal media cells treated with ACTH exhibited more differences than those treated with AngII (Figure 4B). Across all conditions, we detected 3217 genes with significantly different expression levels (FDR <0.05). Consistently, we detected substantially more statistically significant expression changes induced by ACTH versus AngII at 3hrs and 24 hrs (Supplemental Figure 4B, left and right, respectively). However, the expression changes for genes with significant differential expression upon either ACTH or AngII treatment (n=3217) were strongly positively correlated. This indicated that the differences between ACTH and AngII were more quantitative (magnitude of the expression change) rather than qualitative (completely different genes changing) (Figure 4C). Consistent with hormone production, both AngII and ACTH resulted in the induction of genes encoding key steroidogenic enzymes (Figure 4D). Finally we tested if the AngII-induced changes were recapitulated in the ex vivo stimulation paradigm using gene set enrichment analysis (GSEA) (Tamayo *et al*, 2005). Indeed, as2 genes, which were robustly and rapidly induced by AngII in H295R cells, were significantly upregulated after ex vivo treatment of primary adrenocortical cells with ACTH (Figure 4E, top left) and AngII (Figure 4E, bottom left) for 3 hours. Likewise, as4 genes, which also exhibited a robust albeit delayed induction in response to AngII in H295R cells, were significantly upregulated after ex vivo treatment of primary adrenocortical cells with ACTH (Figure 4E, top right) and AngII (Figure 4E, bottom right) for 24 hours. We found 154 RBPs among the union of differentially genes, including ones that were differentially expressed upon AngII treatment of H295R cells, such as CPEB4, MBNL1, MBNL2, MSI2, PEG10, ZFP36, and ZFP36L2. Overall we found strong concordance between the direction and timing of expression changes in H295R cells and primary cells (Supplemental Figure 4C).

**Figure 4.**
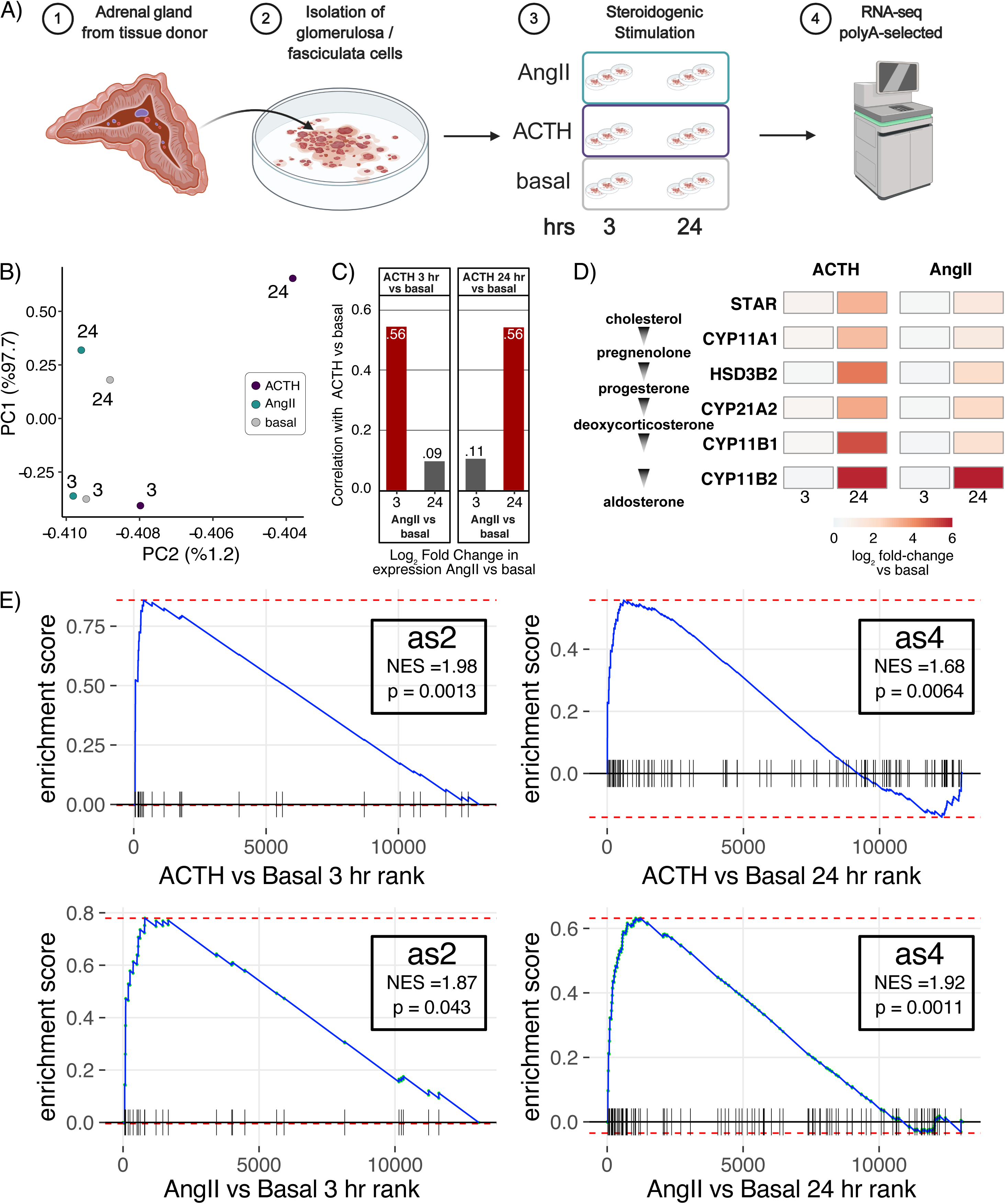
Gene expression dynamics of ex vivo stimulated primary human adrenocortical cells. A) Ex vivo stimulation of cortical cells isolated from normal human adrenal glands treated with AngII, ACTH, or basal media for 3 or 24 hours. Strand-specific paired-end RNA-seq was performed in triplicate on polyadenylated RNA. B) Scatter plot of the first two principal components calculated on a matrix of the mean expression level for each expressed gene at each time (label) and treatment (color). C) Barplot of the Pearson correlation coefficient between AngII-induced and ACTH-induced changes in gene expression versus basal cells for the union of genes exhibiting statistically significant changes in expression in any condition/time. Correlation coefficients for comparisons of the same stimulation time are colored red. D) Heatmap of log2 fold change in expression difference for stimulated (ACTH on left, AngII on right) vs basal for mRNAs encoding key steroidogenic enzymes. F) GSEA running enrichment plots depicting the enrichment of as2 (left) and as4 (right) gene sets representing clusters of genes with similar AngII-induction kinetics in H295R cells (see Figure 1D) in a list of genes ranked by the fold change upon ex vivo stimulation of primary human adrenocortical cells with ACTH (top) or AngII (basal) versus basal media.

### RNA-binding proteins regulate aldosterone levels

Given the agreement in direction and timing of the steroidogenic gene expression response between H295R and primary human cells, we performed an siRNA screen to identify RBPs that regulate aldosterone levels in H295R cells. We selected 16 candidates from an annotated list of 1542 human RBPs (see (Gerstberger *et al*, 2014)) based on 1) differential expression upon AngII treatment in H295R cells; 2) high expression in H295R cells; 3) high and adrenal gland-specific RNA expression across normal human tissues, and 4) RBP motifs enriched in the 3’ UTRs of unstable and destabilized mRNAs (Figure 5A). We used independent siRNAs for each candidate RBP and measured aldosterone levels in cell supernatants 24 hrs after treatment with either vehicle or AngII (see methods for details). Aldosterone levels were normalized by cell viability and compared to a mock electroporation control for both unstimulated and stimulated cell supernatants (Supplemental Figure 5A, Supplemental Table 8). As expected, the depletion of the key steroidogenic transcription factor SF-1 (NR5A1) resulted in loss of aldosterone. Five RBPs exhibited statistically significant increases in aldosterone levels by two independent siRNAs in AngII stimulated cell supernatants (Figure 5B). Among these putative repressors of aldosterone were regulators of global RNA decay (BTG2), ARE-decay (ZFP36L1 and ZFP36L2), and a pseudouridine synthase (TRUB1). We identified 3 potential RBPs for which two independent siRNA knockdowns resulted in decreased aldosterone levels suggesting they promote aldosterone production (Figure 5C). Specifically, these putative activators of aldosterone were a splicing factor (MBNL2), an RNA decay factor (CPEB4), and a signaling scaffold and RNA localization factor (AKAP1). For the seven other RBPs we either observed no change and/or conflicting results for the different siRNAs (HNRNPL, MBNL1, PEG10, PNRC2, ZFP36), or we only screened a single siRNA (MSI2 and PELO) (Figure 5D). Knockdown of RBPs resulted in similar, but more subtle effects on aldosterone levels in unstimulated cells, which was to be expected given the minimal amounts of aldosterone produced in the absence of AngII (Supplemental Figure 5 B-D). These data revealed eight RBPs (10 including RBPs targeted by a single siRNA) that either activated or repressed aldosterone production.

**Figure 5.**
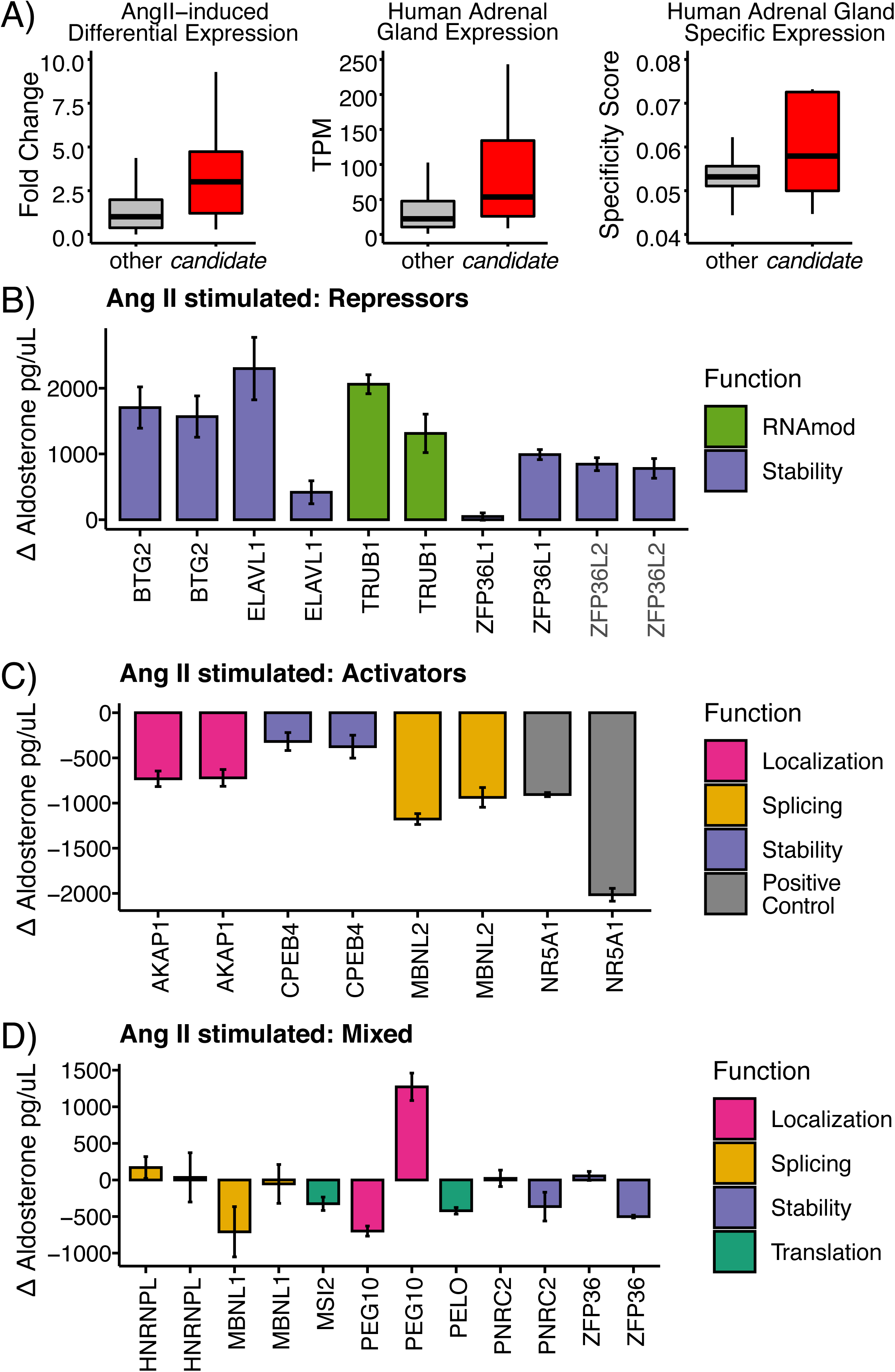
Regulation of aldosterone by RBPs in AngII stimulated cells. A) Boxplots of all RBPs (grey) and candidate RBPs (red) for change in expression upon AngII stimulation (left), human adrenal tissue expression levels (center), and adrenal tissue specificity scores (right). Barplot of the change in aldosterone concentrations for knockdowns resulting in B) increased aldosterone concentration, C) decreased aldosterone concentration, and D) discordant or single siRNA results. Aldosterone levels were measured using ELISA from supernatants of H295R cells electroporated with siRNAs targeting candidate RBPs and stimulated with AngII for 24 hrs. The y-axis represents the change in aldosterone concentration versus mock electroporation (see methods). The error bars represent the standard error of at least 6 replicates. RBPs are color-coded by their known function.

### BTG2 temporally restricts AngII-induced activation kinetics

We decided to investigate how BTG2 controls the AngII gene expression response. BTG2 promotes general RNA decay through deadenylation (Mauxion *et al*, 2008) and represses aldosterone production (Figure 5B). Both BTG2 mRNA (Figure 6A) and protein (Figure 6B) were rapidly induced by AngII stimulation each peaking at ∼3 hours. Therefore, we hypothesize that BTG2 actively promotes the resolution of aldosterone production. We performed RNA-seq of siRNA knockdown of BTG2 in H295R cells that were unstimulated or AngII-stimulated for 6hrs and 24 hr and performed RNA-seq in duplicate (Supplemental Table 9). As expected, BTG2 mRNA levels were lower across all time points in the BTG2 depletion compared to the mock depletion (Figure 6D, right). Next, we assessed the overlap between mRNAs upregulated or downregulated by BTG2 depletion with the AngII-response clusters. We found that mRNAs upregulated upon BTG2 depletion were strongly enriched in the as3, as4 classes and modestly enriched in aw1-aw4 classes (Figure 6C). This is consistent with the propensity for these clusters to contain mRNAs exhibiting increases in RNA decay during steroidogenesis (Figure 3C). Transcripts encoding critical pro-steroidogenic factors and enzymes that were induced by AngII exhibited increased upregulation in the BTG2 depleted cells (Figure 6E), while unchanged mRNAs were not altered by BTG2 (Figure 6D, right). Altogether, these data are consistent with a model in which BTG2 is induced by AngII to upregulate the decay of mRNAs encoding pro-steroidogenic factors to prevent overproduction of aldosterone.

**Figure 6.**
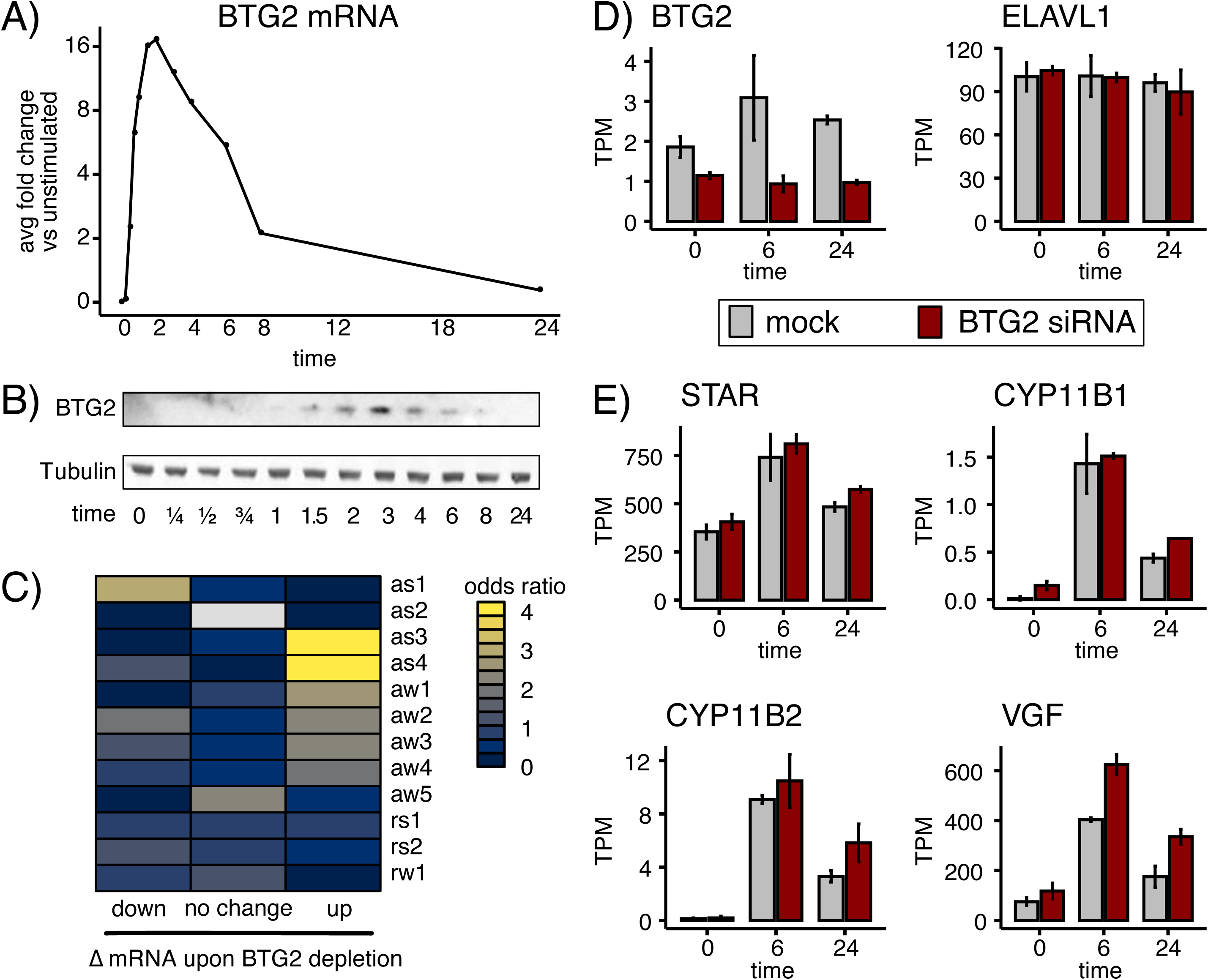
BTG2 constrains AngII RNA expression kinetics to prevent aldosterone overproduction. A) Line plot of fold change in BTG2 mRNA levels in response to AngII. B) Western blot of BTG2 and B-Tubulin protein levels in response to AngII. C) Heatmap of the odds ratio for the overlap between response cluster membership and RNAs exhibiting downregulation, no change, or upregulation upon BTG2 siRNA depletion 24 hours after AngII stimulation (yellow indicates more overlap). D) Barplot of expression levels of BTG2 (downregulated, left) and ELAVL1 (no change, right) upon BTG2 siRNA depletion and AngII stimulation. E) Barplot of expression levels of mRNAs encoding key AngII-induced steroidogenic proteins exhibiting statistically significant upregulation (p<0.05, DESeq2) upon BTG2 siRNA depletion during AngII stimulation. Error bars represent standard deviation.

## DISCUSSION

Despite our current understanding of adrenocortical steroidogenesis, the fundamental regulatory networks that govern aldosterone homeostasis remain unclear. Our data indicate that AngII stimulation increases both mRNA production and decay to rapidly implement and resolve a pro-steroidogenic gene expression program. The AngII-induced expression dynamics in H295R cells are likely to be physiologically-relevant since they were largely recapitulated in primary human adrenocortical cell stimulated with AngII and ACTH. This regulatory scheme enables the robust production of aldosterone while also preventing overproduction. The depletion of multiple factors promoting RNA decay, which themselves were upregulated by AngII, resulted in an excess of aldosterone. The additional energy expenditure of increasing both transcription and decay may represent the cellular cost for the absence of a mechanism to store aldosterone within the cell and release it in response to stimulus. It will be interesting to determine if this coupling is disrupted by mutations driving primary aldosteronism.

### RNA decay temporally coordinates aldosterone production

The stimulation response of adrenocortical cells to AngII resembles an impulse (or single pulse) pattern consistent with functional higher order temporal coordination. This system exhibits both regulatory modules (12 clusters in Figure 1D), which consists of co-expressed genes with the same temporal pattern, which reflects a particular sequential order or cascade of gene or module expression (Yosef & Regev, 2011). The mRNAs in the regulatory modules encoded functionally related proteins that play distinct roles at specific times during the AngII response. For example, expression levels of mRNAs encoding transcription factors and RBPs peaked within the first few hours, while those encoding steroidogenic enzymes peaked 6-8 hours post stimulation. Indeed, the temporal coordination of expression of these distinct factors is consistent with previous characterization of early and late steps of steroidogenesis (Nogueira *et al*, 2009). Our data suggest that both transcriptional cascades and intrinsic differences in RNA decay rates determine the kinetics of the AngII gene expression response. Specifically, the decay rate of transcriptionally induced mRNAs is inversely proportional to the time to peak expression. Therefore, RNA decay is critical for temporally coordinating the response time of these regulatory modules and consequently aldosterone homeostasis. Our findings are similar to the observed role RNA decay in coordinating immune and inflammation responses (Hao & Baltimore, 2009; Elkon *et al*, 2010; Rabani *et al*, 2011). The temporal coordination of these regulatory modules by post-transcriptional regulation may be examples of dynamic RNA regulons (Keene, 2007). Interestingly, these observations occur in systems that require a robust, rapid, yet measured production of a physiologically potent molecule in either cytokines or steroid hormones.

### Coupling of transcription and decay dynamics controls aldosterone homeostasis

Multiple lines of evidence indicate that one or more mixed incoherent feedforward loops (MIFFL) ensure the proper promotion and resolution of steroidogenesis. Specifically, AngII promotes the transcriptional upregulation of both pro-steroidogenic factors (STAR, CYP11B2) and RNA metabolism factors (BTG2, ZFP36L2) that negatively regulate the expression of pro-steroidogenic factors. These Incoherent feedforward loops have largely been examined with respect to transcription factors and miRNAs and are a recurrent motif to enhance the robustness (Tsang *et al*, 2007; Shalgi *et al*, 2007). Transcription factor and RBP versions of these motifs remain poorly characterized (Joshi *et al*, 2012), with the exception of the RBP ZFP36 (TTP) in macrophage activation models (Rabani *et al*, 2014). Nevertheless, the behavioral benefits associated with incoherent feedforward loops include the production of pulse-like patterns, faster response times to stimulation, and even detection of fold-change in expression (Basu *et al*, 2004; Mangan *et al*, 2006; Goentoro *et al*, 2009). These features would be invaluable for facilitating the rapid production and resolution of the response to AngII stimulation to maintain proper aldosterone homeostasis. However due to a combination of technical limitations and need for additional experimentation, we do not know which of those features that BTG2 contributes to. H295R cells are difficult to transfect and it is imperative to develop stable models for controlling gene dose, which we are developing using CRISPR/Cas9. Additionally, we would gain sensitivity by employing recently developed methods that provide a direct readout of newly synthesized vs pre-existing RNAs in the same sample (Herzog *et al*, 2017; Schofield *et al*, 2018). Thus, we are likely underestimating the true number of RNAs exhibiting changes in decay rates during steroidogenesis. Furthermore, we do not know the identity of the transcription factor(s) promoting synthesis of mRNAs encoding pro-steroidogenic proteins and BTG2, though SF1 (NR5A1) and CREB1 are the most likely candidates (Nogueira & Rainey, 2010; Selvaraj *et al*, 2018; Clark & Combs, 1999; Caron *et al*, 1997). The RBPs ZFP36L1 and ZFP36L2 are both induced by AngII and bind to AREs in the 3’ UTR of target mRNA to promote decay and may be additional repressors. Consistent with our expectation, ZFP36L1 was shown to suppress ACTH-stimulated STAR mRNA induction in bovine adrenocortical cells and mouse cell lines (Duan *et al*, 2009). However, this was mediated via ZFP36L1 binding to AREs in an alternative 3’ UTR that we did not detect in our RNA-seq data from H295R or human adrenocortical cells. Finally, BTG2 depletion only deregulated a subset of mRNAs, which suggests that BTG2 may synergize with specific RNA-decay pathways as previously postulated (Stupfler *et al*, 2016).

### Dynamic RNA regulatory network controlling steroidogenesis

We have uncovered a novel paradigm by which RBAs and regulated RNA decay control the kinetics of AngII-stimulated gene expression to facilitate the implementation and resolution of pro-steroidogenic gene expression and metabolic programs required for proper aldosterone production in humans. In addition to RNA decay factors, we identified the RNA modification enzyme TRUB1 was also a repressor of steroidogenesis. TRUB1 is a psuedouridine synthase that modifies tRNAs and mRNAs (Safra *et al*, 2017). We also identified numerous activators of aldosterone production. We found that the signaling scaffold protein AKAP1 activates aldosterone production, which is consistent with a proposed role for AKAP1 in promoting the localized translation of STAR mRNA at the mitochondria (Dyson *et al*, 2008; Grozdanov & Stocco, 2012). Additional activators identified include the cytoplasmic RBPs CPEB4, MSI2, and PELO, which regulated stability and translation, as well as, MBNL2 a well-known splicing factor. The majority of these RBPs have never been associated with aldosterone production or steroidogenesis. The identification of activators and repressors is not surprising given the dysregulation of aldosterone homeostasis has pathological consequences. Dissecting the role of combinatorial regulation by RBPs in the dynamic steroidogenic regulatory network will undoubtedly reveal novel points of control. Since the adrenal cortex is amenable to the delivery of modified oligonucleotides (Biscans *et al*, 2019), the RBP-RNA regulatory interactions may be targeted for disruption to precisely and specifically modulate steroidogenesis.

## METHODS

See supplemental methods for more details.

### Cell culture

H295R cells were cultured in complete media containing DMEM:F12 with 10% Cosmic Calf Serum (Hyclone: SH30087.03) and 1% ITS+ Premix (Corning: 354352). H295R cells were stimulated with 10 nM Angiontensin II (Sigma: A9525) after being in low sera media for 24 hrs (DMEM/F12 with 0.1% CCS, 1% ITS).

Primary human adrenal cells were isolated and cultured as described previously (Xing *et al*, 2010; Rege *et al*, 2015). Primary adrenal cells were plated at a density of 20,000 cells/well (48 well dish) in growth medium and grown to 60% confluence after which they were starved in low serum medium 18 hours prior to treatment with either 10 nM AngII or 10 uM ACTH.

### Gene expression measurements

H295R cells were collected in TRIzol and RNA was isolated using ZYMO Research DirectZol Miniprep Plus kit following the manufacturer’s instructions with on-column DNase I digestion. RNA was collected from *ex vivo* stimulated primary adrenal cells using the Qiagen RNEasy MiniPrep Plus kit following the manufacturer’s instructions and eluted into nuclease-free water.

#### RT-qPCR

Reverse transcription was performed using 100 ng of RNA input for the Bio-Rad iScripit kit and qPCR using Bio-Rad iTaq Universal SYBR Green Supermix on a Bio-Rad CFX 384 qPCR instrument. Analysis of qPCR was performed using the delta-delta Cq method normalizing to GAPDH and unstimulated controls.

#### Metabolic labeling experiments

We performed 4-thiouridine (4sU) experiments as described previously (Mukherjee *et al*, 2017). Briefly, H295R cells were treated for a 20-minute pulse with 500 uM 4sU. RNA was extracted and split into input and biotinylated samples. The biotinylated RNA was immunoprecipitated using streptavidin beads and both input and immunoprecipitated RNA was subject rRNA depletion using RiboZero and sequenced using the qRNA-seq kit (BIOO scientific).

#### RNA-seq and regulatory analysis

Salmon (Patro *et al*, 2017) was used for quantifying transcript levels from all libraries using Gencode v26. A custom salmon index containing Gencode v26 precursor and mature transcripts and ERCC spike-ins as was used previously (Mukherjee *et al*, 2017). All downstream analysis was performed in R and detailed in the Supplemental Methods. Unstimulated (initial) RNA decay rates were estimated using 4sU metabolic labeling data and the INSPEcT R package (de Pretis *et al*, 2015). Overrepresentation of 7mers was calculated via Cwords (Rasmussen *et al*, 2013) from 3’UTR sequences ranked by unstimulated RNA decay rates. RBP motifs enrichment was calculated using https://github.com/TaliaferroLab/FeatureReachR. Determining RNA stability changes during the time course used salmon-quantified intronic and exonic reads, adapting the Exon-Intron Split Analysis concept implemented in eisaR (Gaidatzis *et al*, 2015).

### Aldosterone screen

H295R cells (2.5M) cultured in complete media were electroporated with 10 uM siRNA (Thermo Fisher Scientific - see supplement for catalog numbers) and plated in a 96-well plate. After 24 hrs, half the wells had media was replaced with low sera media and the other half with low sera media containing AngII. After 24hrs, aldosterone levels in the supernatant were measured using Aldosterone Competitive ELISA kit (ThermoFisher) and cell viability measured from the cells using PrestoBlue reagent (ThermoFisher) by reading fluorescence on a BioTek Synergy HT plate reader. Each plate contained a mock transfection that we normalized to for calculating aldosterone levels in each siRNA experiment while taking plate batch effects and cell viability into account.

## DATA ACCESS/SOFTWARE

Raw sequencing data has been deposited in SRA GSE163801. Code used for data analysis is available at https://github.com/mukherjeelab/2020_PTR_steroidogenesis_paper.

## ACKNOWLEDGEMENTS

We would like to thank David Bentley and Lori Sussel for their support, collegiality, and critical review of the manuscript, as well as, Uwe Ohler for initial support for this study. This work was supported by the American Heart Association Predoctoral Fellowship Award 20PRE35220016 (K.W.), the Pre-doctoral Training Grant in Molecular Biology NIH-T32-GM008730 (K.W.), the University of Colorado Anschutz Medical Campus RNA Bioscience Initiative (N.M, R.F.), Boettcher Foundation Webb-Waring Early Career Investigator Award AWD-103075 (N.M), and NIH DK043140 (W.R.).

---

**Supplemental Figure 1.**
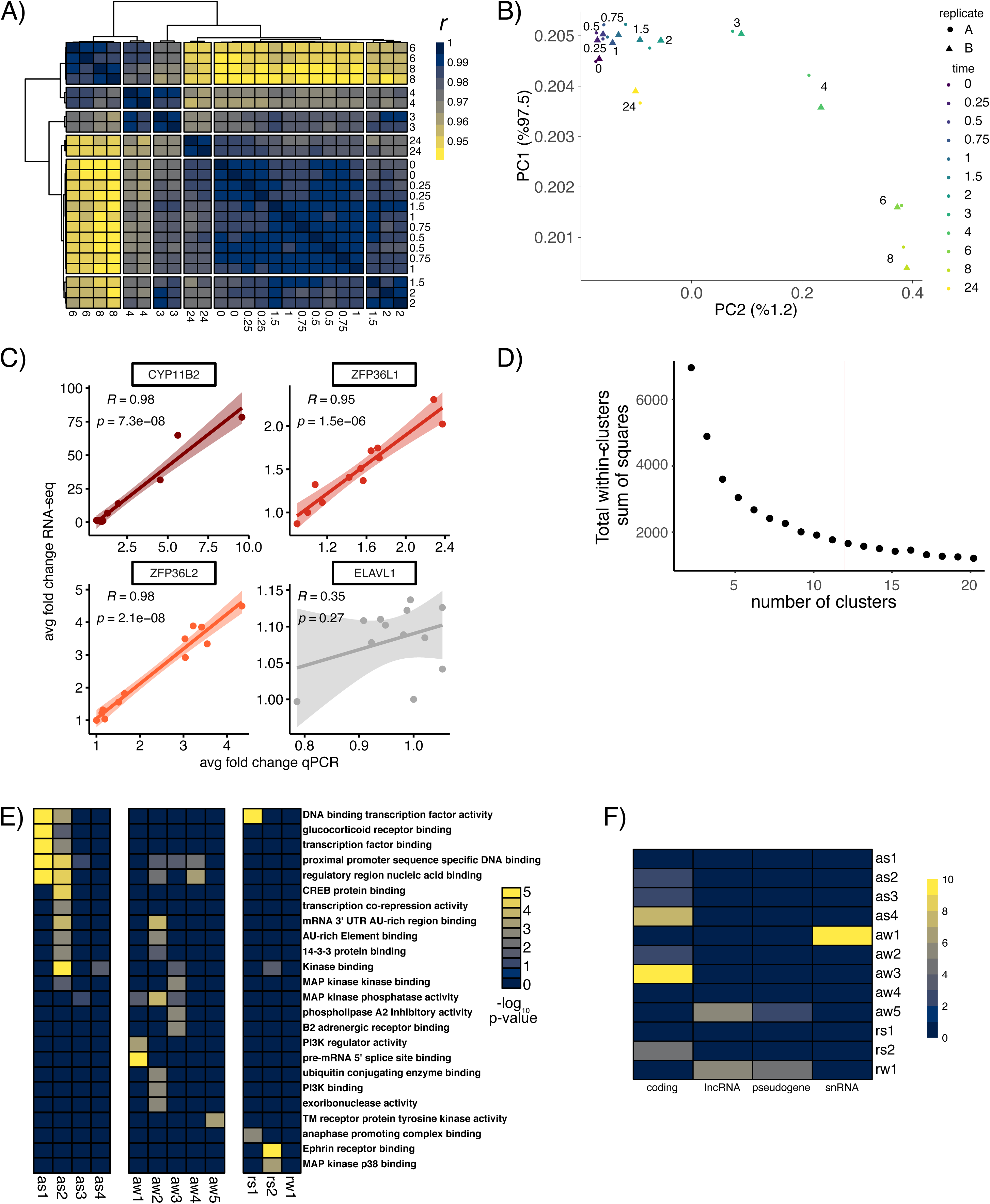
AngII stimulation RNA-seq time course. A) Heatmap of clustered pairwise Pearson correlation coefficients for all samples using expressed genes. B) PCA analysis of all samples using expressed genes. C) Comparison of fold change in expression relative to unstimulated H295R cells between RNA-seq (y-axis) and qRT-PCR. D)Determination of the number of clusters. E) Heatmap of gene biotypes that are significantly enriched in the AngII response groups. F) Heatmap of selected gene sets (Biological Process ontology) that are significantly enriched in the AngII response groups.

**Supplemental Figure 2.**
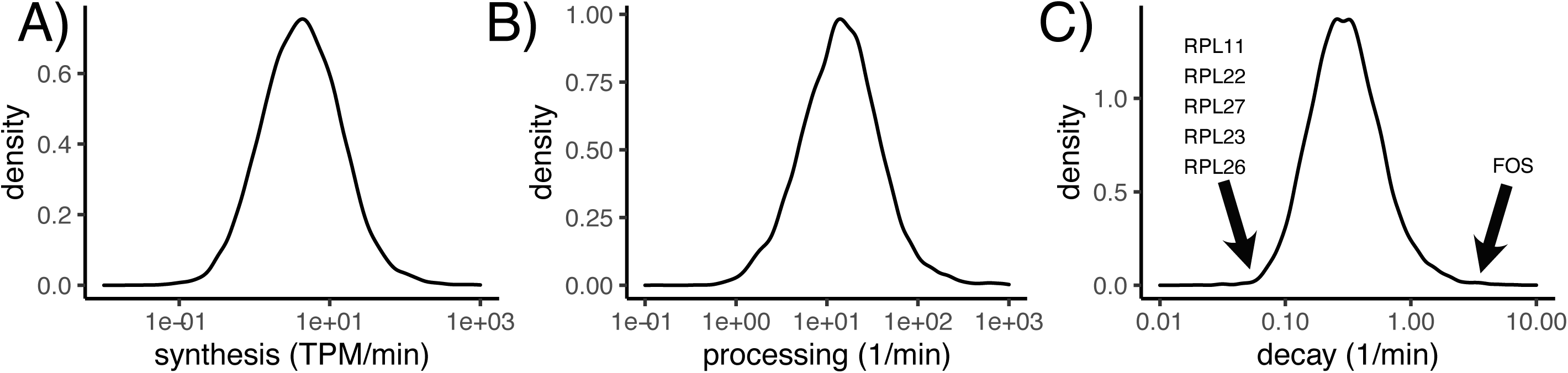
Measuring RNA decay rates in unstimulated H295R cells. Distribution of A) synthesis, B) processing, and C) decay rates calculated using INSPeCT comparing 20-minute 4sU pulse to input RNA. Decay rates of mRNAs encoding ribosomal proteins and immediate early genes depicted.

**Supplemental Figure 3.**
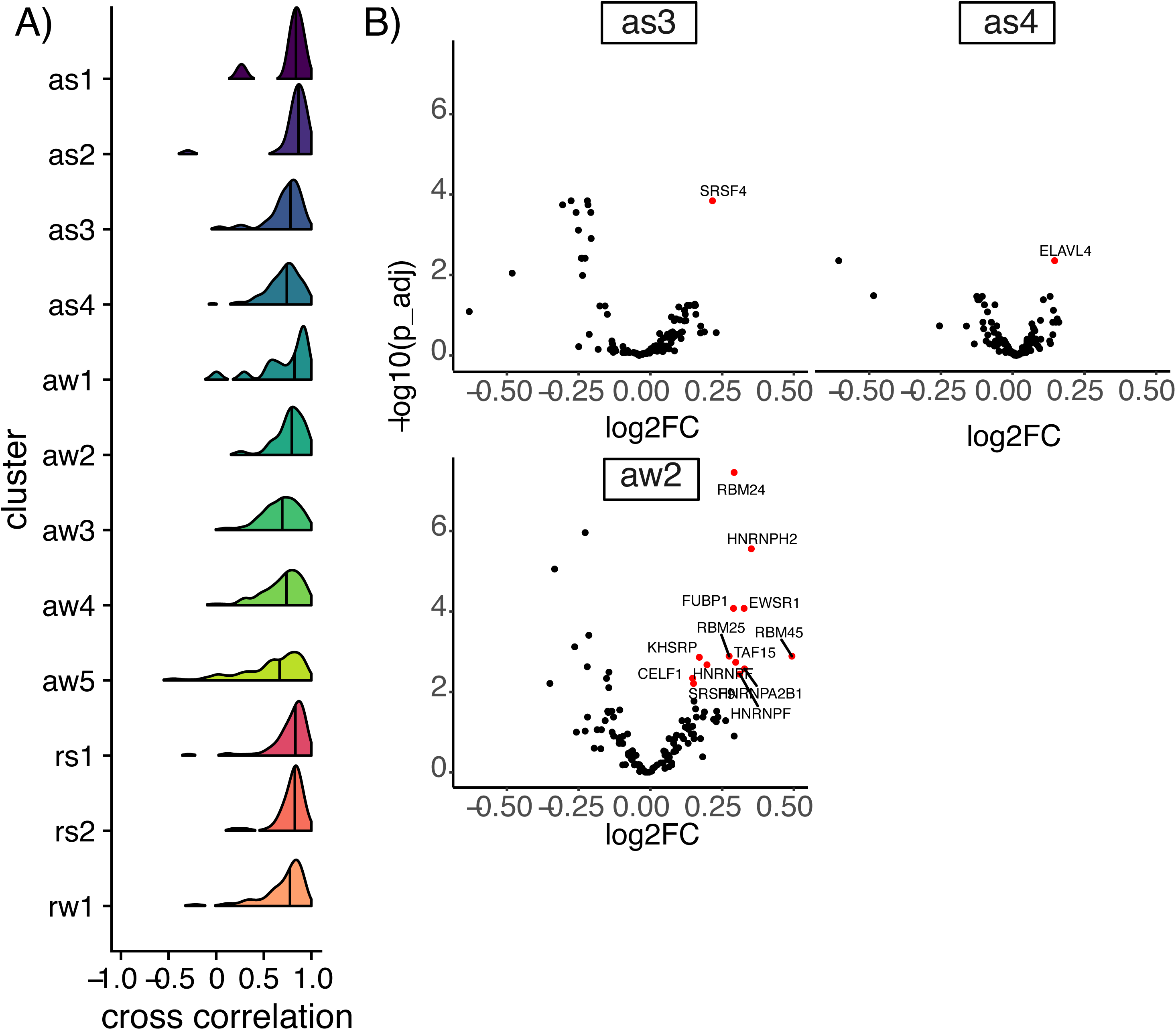
AngII-induced changes in RNA decay. A) Distribution of cross-correlation coefficients between pre-mRNA and mRNA levels for each gene across all time points for each response cluster. B) Scatter plot of the p-value and enrichment of RBP motifs in 5mers within the 3’ UTR of destabilized mRNAs in as3 (top left), as4 (top right), and aw2 (bottom left) versus 3’ UTRs of non-differentially expressed mRNAs.

**Supplemental Figure 4.**
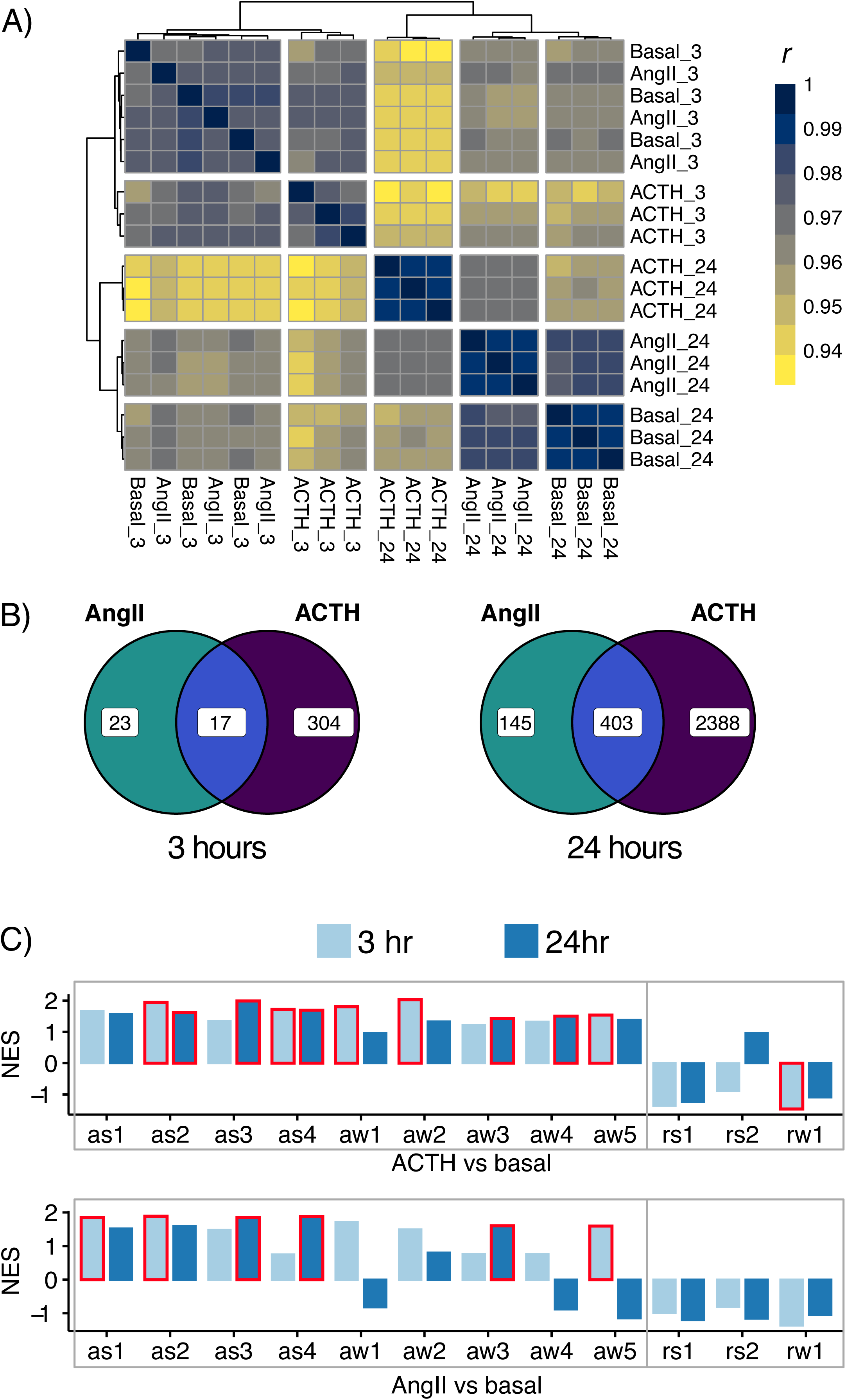
Ex vivo stimulated primary human adrenocortical cells analysis. A) Heatmap of clustered pairwise Pearson correlation coefficients for all samples using expressed genes. B) Venn diagram depicting union and differences between genes differentially expressed upon ex vivo stimulation of primary human adrenocortical cells with AngII or ACTH for 3 hours (left) or 24 hours (right). C) Barplot of normalized enrichment scores (NES) calculated using GSEA representing the enrichment (positive values) or depletion (negative values) of gene sets identified from H295R stimulation response in ACTH (top) or AngII (bottom) induced changes in expression versus basal for each time point. The NES is comparable across gene sets since it adjusts for the size of the gene set and represents the strength of the enrichment/depletion. H295R-derived gene sets with statistically significant enrichment or depletion (p < 0.05) outlined in red.

**Supplemental Figure 5.**
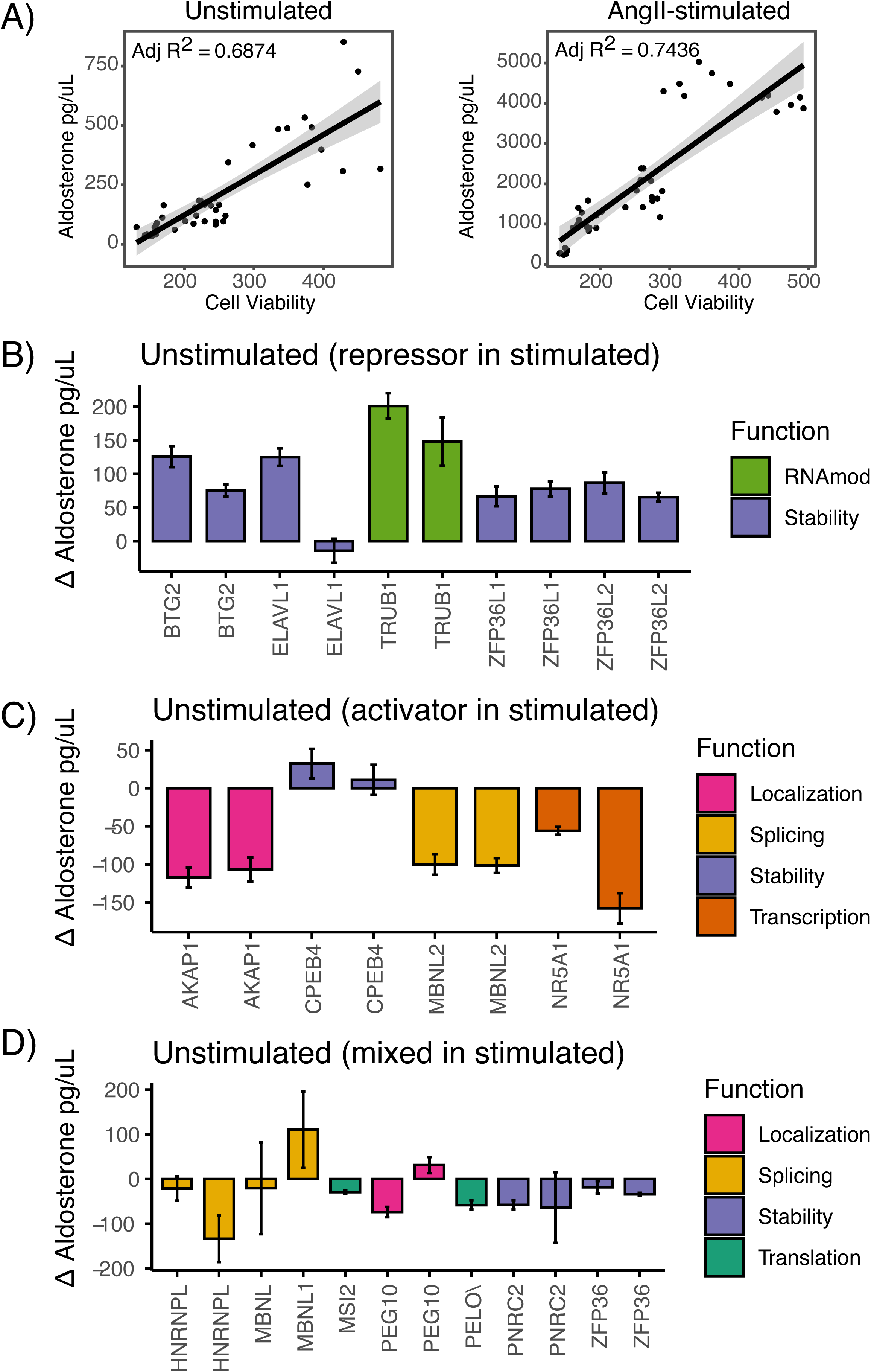
Regulation of aldosterone by RBPs in unstimulated cells. A) Scatterplot of aldosterone levels (supernatant) to cell viability measured using PrestoBlue for unstimulated (left) and AngII-stimulated cells (right). To aid comparison with aldosterone changes from the AngII stimulated cells (Figure 5B-D), the plot arrangement and category assignments (“repressor”, “activator”, “mixed”) are identical to the simulated results. Barplot of the change in aldosterone concentrations for knockdowns resulting in B) increased aldosterone concentration, C) decreased aldosterone concentration, and D) discordant or single siRNA results. Aldosterone levels were measured using ELISA from supernatants of H295R cells electroporated with siRNAs targeting candidate RBPs. The y-axis represents the change in aldosterone concentration versus mock electroporation (see methods). The error bars represent the standard error of at least 6 replicates. RBPs are color-coded by their known function.

## Supplemental Tables

Supplemental Table 1 - AngII-treated H295R mature RNA abundance quantification

Supplemental Table 2 - AngII-treated H295R mature RNA k-means clustering, differential gene expression analysis, and correlation results

Supplemental Table 3 - Estimated decay rates in for unstimulated H295R Supplemental Table 4 - AngII-treated H295R precursor RNA abundance quantification Supplemental Table 5 - Exon-intron-split analysis results

Supplemental Table 6 - AngII-treated adrenocortical cell mature RNA abundance quantification and differential gene expression analysis

Supplemental Table 7 - ACTH-treated adrenocortical cell mature RNA abundance quantification and differential gene expression analysis

Supplemental Table 8 - Changes in aldosterone levels from siRNA screen. Supplemental Table 9 - BTG2 knockdown mature RNA abundance quantification Supplemental Methods - Detailed description of methods used in the manuscript

