## Supplemental Methods for "RNA-binding proteins regulate aldosterone homeostasis in human steroidogenic cells"

**Cell culture**

H295R cells were cultured in complete media containing DMEM:F12 with 10% Cosmic Calf Serum (Hyclone: SH30087.03) and 1% ITS+ Premix (Corning: 354352). All experiments involving the stimulation of H295R cells with Angiotensin II were performed in the following manner. On the first day, H295R cells were plated in complete media. On the second day, cells were switched to low sera media (DMEM/F12 with 0.1% CCS, 1% ITS). On the third (24 hr) or fourth-day cells were treated with 10 nM AngII at different times and simultaneous collection on day 4. 500,000 cells were plated on day 1 and performed in duplicate for data collected from 24-hour time course of H295R cells stimulated with AngII.

Primary human adrenal cells were isolated and cultured as described previously (Rege *et al*, 2015; Bassett *et al*, 2004). Briefly, adrenal tissue was minced and dissociated into a single-cell suspension by repeated exposure of the tissue fragments to DMEM/F12 medium containing 1 mg/mL collagenase dispase and 0.25 mg/mL DNaseI (Hoffmann-La Roche Ltd). Four 1-hour cycles of digestion at 37°C and mechanical dispersion were performed. Cells were collected between each digestion and combined prior to storage at −150°C. Primary adrenal cells were plated at a density of 20,000 cells/well (48 well dish) in growth medium and grown to 60% confluence after which they were starved in low serum medium 18 hours prior to treatment with either 10 nM AngII or 10 uM ACTH.

**RNA sequencing**

For H295R cells stimulated with human Angiotensin II across 24-hour time course, RNA was collected by removing media, washing cells briefly in 1X PBS, followed by the addition of TRIzol Reagent directly to the wells containing the cells. The cells were then incubated at room temperature for 5 minutes followed by scraping the wells with a cell lifter. The samples in TRIzol were collected into new tubes and frozen at -80°C. The RNA was later isolated using the ZYMO Research DirectZol Miniprep Plus kit following the manufacturer’s instructions with on-column DNase I digestion. RNA was eluted from the column with nuclease-free water and quantified by Nanodrop and Qubit RNA BR assay. cDNA and minus reverse transcriptase control samples were generated for each sample from 100ng total RNA input using Bio-Rad iScript kit following the manufacturer’s direction, including a water no-template control sample for reverse transcriptase-containing reactions. The resulting reactions were diluted with nuclease-free water and 2 μL used as input into 10 μL qPCR reaction with a final concentration of 1X Bio-Rad iTaq Universal SYBR Green Supermix and 500nM primers. qPCR reactions were performed in triplicate for each sample and primer pair combination, including minus RT and no-template RT control reactions. Primer pairs amplified regions of GAPDH, ELAVL1, CYP11B2, ZFP36L1, and ZFP36L2 genes and are listed at the bottom of the document. Reactions were run on a Bio-Rad CFX 384 qPCR instrument with parameters as described in the manufacturer’s instructions for the Bio-Rad iTaq SYBR Green supermix. 500ng total RNA was used as input into ribosomal RNA depletion followed by RNA-seq library preparation using KAPA Biosystems KAPA RNA Hyper Prep kit with RiboErase and dual index adapters. Final libraries were amplified with 10 cycles of PCR and sequenced on the NovaSeq 6000 for 2x150 bp PE reads at the University of Colorado School of Medicine Genomics and Microarray Core Facility.

RNA was collected from *ex vivo* stimulated primary adrenal cells using the Qiagen RNEasy MiniPrep Plus kit following the manufacturer’s instructions and eluted into nuclease-free water. RNA samples were quantified by Nanodrop. PolyA containing RNA was collected for each sample from approximately 65ng total RNA input using one round of the Lexogen PolyA selection kit following the manufacturer’s instructions. PolyA selected RNA was eluted in nuclease-free water and used as input in downstream RNA-seq library preparation using the KAPA Biosystems KAPA RNA Hyper Prep kit and dual index adapters. Final libraries were amplified with 15 cycles of PCR and sequenced on the NovaSeq 6000 for 2x150 bp PE reads at the University of Colorado School of Medicine Genomics and Microarray Core Facility. An aliquot of each final library sample was diluted with nuclease-free water to 0.5 ng/μL and 2 μL used as input into 10 μL qPCR reaction with a final concentration of 1X Bio-Rad iTaq Universal SYBR Green Supermix and 500 nM primers. qPCR reactions were performed in triplicate for each sample and primer pair combination, including minus RT and no-template RT control reactions. Primer pairs amplified regions of GAPDH, ELAVL1, CYP11B2, and ZFP36L2 genes and are listed at the bottom of this document. Reactions were run on a Bio-Rad CFX 384 qPCR instrument with parameters as described in the manufacturer’s instructions for the Bio-Rad iTaq SYBR Green supermix. Analysis of qPCR was performed using the delta-delta Cq method normalizing to GAPDH and unstimulated controls.

**Data analysis**

The code used for data analysis is available [here](https://github.com/mukherjeelab/2020_PTR_steroidogenesis_paper). Read mapping and quantification via salmon (Patro *et al*, 2017) used settings “-l A --allowDovetail –validateMappings” and a custom salmon index containing Gencode v26 precursor and mature transcripts and ERCC spike-ins as was used previously (Mukherjee *et al*, 2017). Next, salmon quantification tables were imported via tximport into edgeR framework for differential expression analysis using generalized linear models. Differentially expressed genes, at FDR < 0.001 cutoff, were then grouped by k-means clustering, with the number of clusters optimized by the elbow method. Line plots for each cluster summarized the average fold change and standard error of the mean.

GeneOverlap and msigdbr R packages provided Gene Ontology Biological Process term enrichment for the clusters. Expression profile “peakiness”, measurement of the sharpness of peaks or maximal peak height, was calculated according to the previously described formula (Rabani *et al*, 2011). Calculations of cross-correlation and lag values between precursor and mature RNA profiles of each gene was achieved with custom code and base R functions. Unstimulated (initial) RNA degradation rates were estimated using 4sU metabolic labeling data and the INSPEcT R package (de Pretis *et al*, 2015). Overrepresentation of 7mers was calculated via Cwords (Rasmussen *et al*, 2013)and 3’UTR sequences ranked by unstimulated RNA degradation rates. The top 500 7mers were then processed through FeatureReachR to report associated enriched RNA-binding protein annotations.

Modeling of RNA stability changes during the time course used salmon-quantified intronic and exonic reads, adapting the Exon-Intron Split Analysis concept implemented in eisaR. The odds ratio for overlap between clusters and EISA-determined RNA stability change was obtained via GeneOverlap. Enriched 3’UTR motifs in cluster “aw3” destabilized genes were calculated with position weight matrix scanning of RBNS motif annotations, using all 3’UTRs of genes with no differential expression (edgeR) and no change in RNA stability (eisaR) within the AngII time course as control sequence set, via FeatureReachR (https://github.com/TaliaferroLab/FeatureReachR).

**Aldosterone screen**

H295R cells were grown to approximately 80% confluency in complete media [DMEM:F12 with 10% Cosmic Calf Serum (Hyclone: SH30087.03) and 1% ITS+ Premix (Corning: 354352)], washed with 1X PBS (without Ca^+2^ and Mg^+2^), dissociated with TrypLE, then collected into complete media on day 1. An aliquot of the resulting single-cell suspension was counted using the Invitrogen Countess. Prior to electroporation, 2,475,000 cells from the single-cell suspension were centrifuged at 400g for 5 minutes, media removed, and the resulting cell pellet washed with room temperature 1X PBS. The washed pellet was resuspended in 110 μL “R buffer” provided in the Invitrogen Neon 100 μL electroporation kit (ThermoFisher Scientific: MPK10096). 100 μL of the cells in R buffer were further mixed with 12.5 μL of 10uM siRNA of interest (purchased from Thermo Fisher Scientific, siRNAs listed below) or water for mock electroporation (water mock electroporation was included for each plate) 100 μL of the siRNA (or water mock) and cell suspension mixture (final cell count equaling 2 million) was aspirated using the Invitrogen Neon pipette fitted with a 100 μL electroporation tip and electroporated in 3 mL E2 buffer in the kit-provided acrylic tubes with the following settings: Voltage, 1100v; Width, 30ms; Pulses, 2. After electroporation, the cells were dispensed into a conical tube containing 7.9 mL of pre-warmed complete H295R media and gently mixed. The electroporated cells in complete media were kept at 37°C, 5% CO_2_ until the remaining siRNAs or water mock control electroporations were performed. For each experimental batch, 5 siRNAs and 1 water mock electroporation were performed. When possible, two unique siRNAs for each gene of interest were used separately for electroporation. Upon completion of the electroporations, the cells in each tube were resuspended by pipetting, transferred to a sterile reagent reservoir, and 100 μL of cell suspension (25,000 cells) were transferred to 12 wells (one row) in a 96-well cell culture plate for later supernatant collection for ELISA. The remaining 2 rows of the 96-well plate were filled with 100uL of complete media in each well. 5 mL (1,250,000 cels) of the remaining electroporated cell suspension was plated into one well of a 6 well plate for later RNA collection. The plates were incubated at 37°C, 5% CO_2_ for 24-25 hours.

On day 2 after 24-25 hour incubation, media was aspirated from all wells and cells were gently washed with pre-warmed 1X PBS (without Ca^+2^ and Mg^+2^) then aspirated. For each row of electroporated cells in the 96-well plate, 6 of the 12 wells were replaced with 110 μL low-sera media (DMEM/F12 with 0.1% CCS, 1% ITS) or low-sera media containing a final concentration of 10 nM Angiotensin II (AngII). Media-only wells were replaced with low-sera media not containing AngII. Media in the 6-well plate was removed, cells washed gently with 1X PBS (without Ca^+2^ and Mg^+2^), and replaced with low-sera media containing 10 nM AngII. The plates were incubated at 37°C, 5% CO_2_ for 24 hours.

On day 3 after 24 hour incubation, media was aspirated from each well of the 6-well plate, cells gently washed with pre-warmed 1X PBS (without Ca^+2^ and Mg^+2^), and cells collected into 600 μL RNA Lysis Buffer from the ZYMO Research Quick RNA Micro kit. Cell lysate was stored at -80°C for future use. 100 μL of supernatant from each well of the 96-well plate was removed and transferred to the appropriate well of the Invitrogen Aldosterone Competitive ELISA kit (ThermoFisher: EIAALD) plate (see below). The resulting plate containing adhered cells was aspirated to remove any residual media and 100 μL 1X PrestoBlue reagent (ThermoFisher: ) made with H295R complete media was added to each well and incubated at 37°C for 30 minutes. During incubation, ELISA was performed as outlined below. PrestoBlue plate was read on a BioTek Synergy HT plate reader after 30 minutes incubation and again the next day after 24 hour incubation. Each ELISA plate contained 5 siRNA electroporations and 1 water mock electroporation (each consisting of 6 wells of 24 hour AngII 10 nM stimulated cells and 6 wells unstimulated cells in low-sera media), 8 wells of low-sera media only control, 7 standards and “NSB” controls as provided and directed by the manufacturer. The Aldosterone competitive ELISAs were performed as instructed by the manufacturer and assay read on a BioTek Synergy HT plate reader.

Since the number of cells plated per electroporation could influence the amount of aldosterone we detected, we included a mock transfection on each plate. Specifically, we constructed a multiple regression model using the plate (batch) and PrestoBlue on the mock-transfected samples to estimate aldosterone levels. We then collected the residuals from the regression, which represent the variance in aldosterone levels not explained by batch or cell number/viability. We plotted the residuals for each siRNA to visualize the difference in aldosterone levels per experiment. Statistical significance was calculated using ANOVA.

| GeneID | AssayID |
| --- | --- |
| MSI2 | s42756 |
| NR5A1 | s5393 |
| NR5A1 | s5392 |
| ZFP36 | s14979 |
| ZFP36 | s14978 |
| CPEB4 | s37202 |
| CPEB4 | s37201 |
| ELAVL1 | s4610 |
| BTG2 | s15386 |
| BTG2 | s15385 |
| PELO | s28808 |
| PEG10 | s23004 |
| PEG10 | s23005 |
| TRUB1 | s44502 |
| TRUB1 | s44503 |
| MBNL2 | s19767 |
| MBNL2 | s19768 |
| AKAP1 | s15665 |
| AKAP1 | s15666 |
| PNRC2 | s31122 |
| MBNL1 | s8553 |
| MBNL1 | s8555 |
| HNRNPL | s6740 |
| HNRNPL | s6741 |
| PNRC2 | s31123 |
| Scramble Negative Control #1 | AM4611 |

| qPCR Oligos |  |
| --- | --- |
| Name | Sequence |
| GAPDH_hsa_qFWD | CAT TGC CCT CAA CGA CCA CTT TGT |
| GAPDH_hsa_qREV | TCT ACA TGG CAA CTG TGA GGA GGG |
| ELAVL1_hsa_qFWD | CGC CAA CTT GTA CAT CAG CG |
| ELAVL1_hsa_qREV | TAA ACG CAA CCC CTC TGG AC |
| CYP11B2_hsa_qFP1 | GGC AGA GGC AGA GAT GCT G |
| CYP11B2_hsa_qRP1 | CTT GAG TTA GTG TCT CCA CCA GGA |
| ZFP36L1_hsa_qFWD | GTC TGC CAC CAT CTT CGA CT |
| ZFP36L1_hsa_qREV | TTT CTG TCC AGC AGG CAA CC |
| ZFP36L2_hsa_qFWD | CAC TGC GGG ATC CAG AAA CA |
| ZFP36L2_hsa_qREV | TGA GGT TGG CCA GGG ATT TC |

Bassett MH, Suzuki T, Sasano H, White PC & Rainey WE (2004) The orphan nuclear receptors NURR1 and NGFIB regulate adrenal aldosterone production. *Mol Endocrinol* 18: 279–290

Mukherjee N, Calviello L, Hirsekorn A, de Pretis S, Pelizzola M & Ohler U (2017) Integrative classification of human coding and noncoding genes through RNA metabolism profiles. *Nat Struct Mol Biol* 24: 86–96

Patro R, Duggal G, Love MI, Irizarry RA & Kingsford C (2017) Salmon provides fast and bias-aware quantification of transcript expression. *Nat Methods* 14: 417–419

de Pretis S, Kress T, Morelli MJ, Melloni GEM, Riva L, Amati B & Pelizzola M (2015) INSPEcT: a computational tool to infer mRNA synthesis, processing and degradation dynamics from RNA- and 4sU-seq time course experiments. *Bioinformatics* 31: 2829–2835

Rabani M, Levin JZ, Fan L, Adiconis X, Raychowdhury R, Garber M, Gnirke A, Nusbaum C, Hacohen N, Friedman N, *et al* (2011) Metabolic labeling of RNA uncovers principles of RNA production and degradation dynamics in mammalian cells. *Nat Biotechnol* 29: 436–442

Rasmussen SH, Jacobsen A & Krogh A (2013) cWords - systematic microRNA regulatory motif discovery from mRNA expression data. *Silence* 4: 2

Rege J, Nishimoto HK, Nishimoto K, Rodgers RJ, Auchus RJ & Rainey WE (2015) Bone Morphogenetic Protein-4 (BMP4): A Paracrine Regulator of Human Adrenal C19 Steroid Synthesis. *Endocrinology* 156: 2530–2540
